# Detection of Parkinson’s disease using blood and brain cells transcript profiles

**DOI:** 10.1101/483016

**Authors:** Mohammad Ali Moni, Humayan Kabir Rana, M. Babul Islam, Mohammad Boshir Ahmed, Md. Al Mehedi Hasan, Fazlul Huq, Julian M.W. Quinn

## Abstract

Identification of genes whose regulation of expression is similar in both brain and blood cells could enable monitoring of significant neurological traits and disorders by analysis of blood samples. We thus employed transcriptional analysis of pathologically affected tissues, using agnostic approaches to identify overlapping gene functions and integrating this transcriptomic information with expression quantitative trait loci (eQTL) data. Here, we estimate the correlation of genetic expression in the top-associated cis-eQTLs of brain tissue and blood cells in Parkinson’s (PD).

We introduced quantitative frameworks to reveal the complex relationship of various biasing genetic factors in PD, a neurodegenerative disease. We examined gene expression microarray and RNA-Seq datasets from human brain and blood tissues from PD-affected and control individuals. Differentially expressed genes (DEG) were identified for both brain and blood cells to determine common DEG overlaps. Based on neighborhood-based bench-marking and multilayer network topology aproaches we then developed genetic associations of factors with PD.

Overlapping DEG sets underwent gene enrichment using pathway analysis and gene ontology methods, which identified candidate common genes and pathways. We identified 12 significantly dysregulated genes shared by brain and blood cells, which were validated using dbGaP (gene SNP-disease linkage) database for gold-standard benchmarking of their significance in disease processes. Ontological and pathway analyses identified significant gene ontology and molecular pathways that indicate PD progression.

In sum, we found possible novel links between pathological processes in brain and blood cells by examining cell pathway commonalities, corroborating these associations using well validated datasets. This demonstrates that for brain-related pathologies combining gene expression analysis and blood cell cis-eQTL is a potentially powerful analytical approach. Thus, our methodologies facilitate data-driven approaches that can advance knowledge of disease mechanisms and may enable prediction of neurological dysfunction using blood cell transcript profiling.

## 1. Introduction

The difficulty of early diagnosis and discrimination of neurodegenerative diseases such as Parkinson’s (PD), Alzheimer’s (AD), Huntington’s (HD) and motor neuron diseases (MND) mean that identifying robust biomarkers for these conditions in blood samples is a long held goal [1, 2, 3]. While gene expression studies of peripheral blood mononuclear cells cannot directly study the diseased central nervous tissues at issue, it is an approach with the potential to characterize the influences of systemic factors that similarly affect both brain and blood cells. These influences includes genetic factors, notably protein coding-gene mutations and single nucleotide polymorphisms (SNPs), as well as other systemic diseases (i.e., comorbidities) that are suffered by neurodegenerative disease patients. The latter can be an important consideration since many comorbidities are known risk factors for neurodegenerative diseases [4, 5, 6]. However, since most genes are regulated differently in different tissues, to obtain useful biomarkers among blood cell transcripts we need to identify genes that are similarly regulated in brain and blood cells. Achieving this could enable blood cell transcript profiles to become a window for viewing some of the pathological changes affecting the brain.

Genome-wide association studies (GWAS) have discovered thousands of genetic variants associated with complex traits and pathological conditions, including neurodegenerative diseases [7, 8]. With advances in microarray and RNA-Seq technologies, genome-wide sequencing together with tissue gene expression data from relatively large numbers of samples have been generated as part of attempts to identify genetic variants that affect transcript abundance [9, 10], i.e., expression quantitative trait loci (eQTLs). The Genotype-Tissue Expression (GTEx) project [11] has provided a large and increasingly comprehensive resource of data that enable investigation of the genetic causes of gene expression variation across a broad range of tissues and cell types, particularly including blood cells and brain tissue. Molecular associations, evident from differential gene expression patterns, proteinprotein interactions (PPIs), gene ontologies and common metabolic pathways can mediate the effects risk factors that influence or drive development of a disease [12, 13, 14]. A risk factor (e.g., a pathogen or a comorbidity) and a disease may reveal a mechanistic link if both cause altered expression in a common set of genes [15, 16, 17]. In addition, from a proteomics and signaling pathways perpective, such links may also be found through the demonstrations of common biological modules such as PPIs, gene ontologies or molecular pathways [18, 19, 20]. We thus used a network based analyses to identify the influence of genetic factors and disorders in PD progression by utilizing the gene expression profiling, PPI sub-network, gene ontologies and molecular pathways. An extensive study regarding phylogenetic and pathway analysis was also conducted to reveal the genetic associations of the PD.

In this study, we have investigated PD, a progressively developing degenerative disorder that mainly damages the motor system of the central nervous system [21] suffered by over 10 million people worldwide [22]. PD damages dopaminergic neurons in the substantia nigra pars compacta and forms Lewy bodies, neuronal cell soma inclusions containing *α*-synuclein [23]. Subtle early symptoms seen in PD affected individuals comprise shaking, rigidity, slowness of movement and complications in mobility. PD patients suffer from difficulties in walking, talking or even completing simple daily activities, while sensory, sleep and emotional problems may also be evident, and can lead to development of dementia [24]. Currently the main causes or risk factors of PD remain poorly understood [25]. For example, it unclear why the risk of PD development is affected by gender or by physical exercise [26]. To gain mechanistic insight into PD (and potentially other central nervous system disorders) we investigated PD-associated transcripts (from RNA-Seq and microarray data) that are common to brain and blood cells, then used human-expressed quantitative trait loci (eQTL) data to identify biomarkers that are expressed under similar genetic control in both cell types. Further screens and filtering methods that employ human genetics and transcriptomics databases (including microarray and RNAseq data), as well as curated gold standard benchmark databases for PD was used to reveal potential brain biomarker genes and cell pathways which behave similarly in blood. Our overall approach aims to find overlapping pathways of potential clinical utility, but with the possibility that we can also identify important new pathways relevance to many neurological diseases by examining blood cell transcripts.

## 2. Materials and Methods

### 2.1 Overview of analytical approach

We present here, sumarised in Fig. 1, a systematic and quantitative approach to using blood cell information to predict neurological disorder at the early stages using available sources of mRNA expression, RNA-Seq, Genome Wide Association Studies (GWAS) and expression of quantitative trait loci (eQTL) datasets. This approach employs gene expression analyses, signalling pathway information, Gene Ontology (GO) data, disease–gene associations and protein-protein interaction data to identify putative components of common pathways for the neurological dysfunction evident in brain and blood cells.

**Figure 1:**
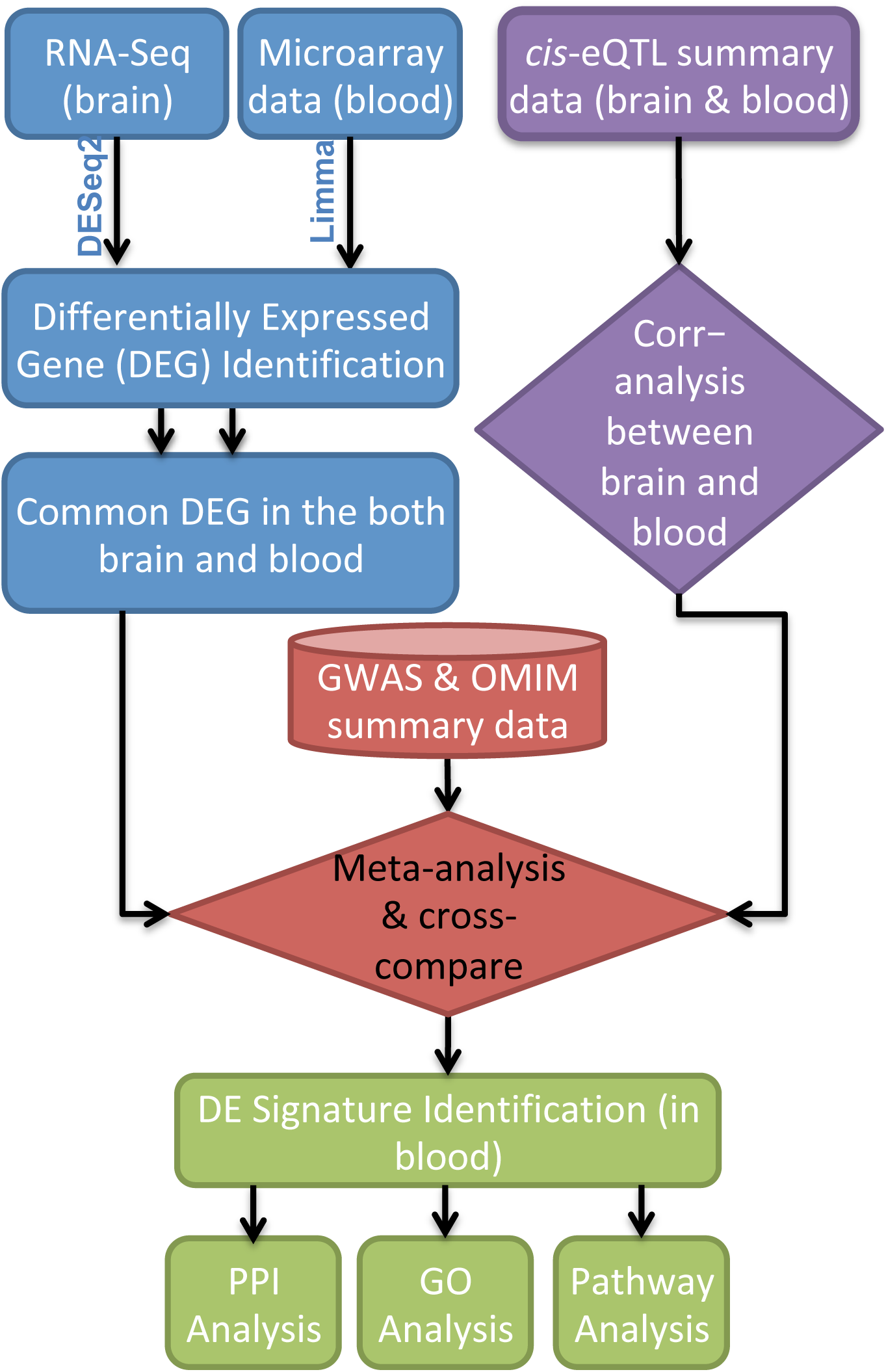
Flowchart of the pipeline that is used for the early detection of the neurological dysfunction using blood cell transcripts.

### 2.2 Datasets employed in this study

To investigate the molecular pathways involved in PD, we first employed global transcriptome analyses (RNAseq datasets) as well as gene expression microarray datasets related to the blood and brain cells. In this study, we collected raw data from the Gene Expression Omnibus of the National Center for Biotechnology Information (NCBI) (http://www.ncbi.nlm.nih.gov/geo/). We selected 2 different large human gene expression datasets for our study with accession numbers GSE68719 and GSE22491. GSE68719 is an RNAseq dataset from a study of PD using brain cells from healthy and PD individuals. GSE22491 is an Affymetrix RNA array dataset from a study of normal and PD individuals whole blood cells. We have also used GWAS catalogues is the NHGRI-EBI Catalog of published genome-wide association studies (https://www.ebi.ac.uk/gwas/) and eQTL data (both blood and brain) from the GTExPortal which is a database for the Genetic Association data (https://gtexportal.org/home/)

To get further insight into the molecular pathways of PD that overlap between brain and blood cells, we performed pathway and gene ontology analysis using the DAVID bioinformatics resources (https://david-d.ncifcrf.gov/) and KEGG pathways database [27]. We also generated a protein-protein interaction (PPI) network for each disease-pair datasets, using data from the STRING database string-db.org citeamberger2017searching. Furthermore, we also incorporated a gold bench mark verified dataset dbGaP (www.ncbi.nlm.nih.gov/gap) in our study for validating the proof of principle of our network based approach.

### 2.3 Analysis methods

Using RNAseq and RNA microarray technologies for global transcriptome analyses, we compared the gene expression profiles of PD with that in brain and blood cells datasets. All these datasets were generated by comparing diseased tissue with normal to identify differentially expressed genes (DEG) associated with their respective pathology PD. The analysis was performed from the original raw datasets and employed DESeq2 [28] and Limma [29] R Bioconductor packages for the RNAseq and microarray data respectively. To avoid the problems of comparing mRNA expression data of different platforms and experimental systems, we normalized and calibrated the gene expression data in each sample (disease state or control) using the Z-score transformation (*Z*_*ij*_) for each disease gene expression matrix using:

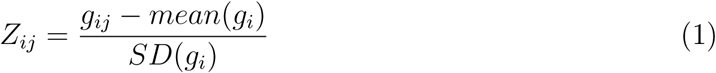

where *SD* is the standard deviation, *g*_*ij*_ represents the expression value of gene *i* in sample *j*. This transformation allows direct comparisons of gene expression values across samples and diseases. We applied a Studentised t-test statistic between two conditions. Data were *log*_2_-transformed and Student’s unpaired t-test performed to identify genes that were differentially expressed in patients over normal samples and significant genes were selected. A threshold of at least 1 *log*_2_ fold change and a *p*-value for the t-tests of < 5 *×* 10^*-*2^ were chosen. In addition, a two-way ANOVA with Bonferroni’s *post hoc* test was used to establish statistical significance between groups (< 0.01).

For gene-disease associations, we applied neighborhood-based benchmarking and topological methods. Given a particular set of human diseases *D* and a set of human genes *G*, gene-disease associations attempt to find whether gene *g* ∈ *G* is associated with disease *d* ∈ *D*. If *G*_*i*_ and *G*_*j*_, the sets of significant up- and down-dysregulated genes associated with diseases *i* and *j* respectively, then the number of shared dysregulated genes 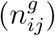 associated with both diseases *i* and *j* is as follows:

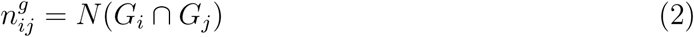

The common neighbours identified by an approach based on the Jaccard Coefficient method [30, 31], where the edge prediction score for the node pair is:

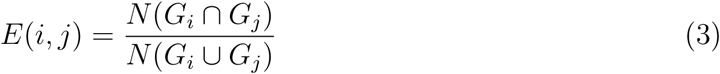

where *E* is the set of all edges. We have used our own R software packages “comoR” [32] and “POGO” [33] to compute novel estimators of the disease comorbidity associations.

### 2.4 Association of eQTL effects between tissues

Let *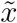* be the estimated effect at the top-linked cis-eQTL for a gene. We can calculate *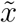* as

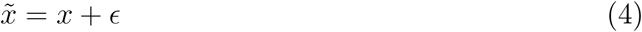

where *x* is the true effect and *ϵ* is the estimated error. We assume that *x* and *ϵ* are random variables across the genes, i.e., *x ∼ N* (0, *var*(*x*)) and *epsilon ∼ N* (0, *var*(*e*)). The covariance of the estimated cis-eQTL effects between tissues *i* and *j* across genes can be partitioned into the co-variance of true cis-eQTL effects and the co-variance of estimation errors (if sample overlap exist), i.e.,

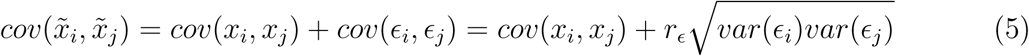

where *var*(ϵ_*i*_) and *var*(ϵ_*j*_) are the variance of the estimated errors across genes in tissues *i* and *j* respectively, and *r*_*ϵ*_ is the correlation of estimated errors across genes between two tissues, i.e., *r*_*ϵ*_ = *cor*(*ϵ*_*i*_, *ϵ*_*j*_). We know that *r*_*ϵ*_*≈ r*_*p*_*ρ*, where 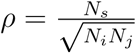 measures the sample overlap with *N − i* and *N − j* being the sample sizes in tissues *i* and *j*, respectively, and *N*_*s*_ being the number of overlapping individuals, and *r*_*p*_ is the association of gene expression levels between two tissues in the overlapping sample. If *i* = *j*, then *r*_*ϵ*_ = 1 and 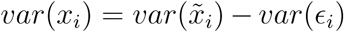, where *var*(*x*_*i*_) is the variance of true cis-eQTL effects across genes in tissue *i*. Thus We can estimate the correlation of true cis-eQTL effect sizes across genes between tissues *i* and *j* as

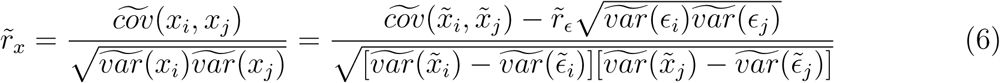

where 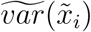 and 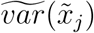 are the observed sample variances of *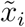* and *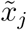*, respectively, in a set of genes, and 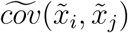 is the observed sample covariance between *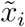* and *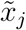* the set of genes.

From Eq. (5) we know that if 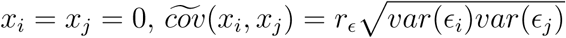. Hence, 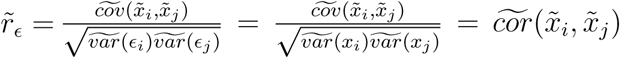 for null SNPs, where 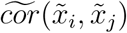 the observed sample association between *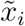* and 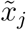 the set of genes. In practice, we calculated 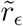 for each gene using null SNPs (PeQTL*>* 0.01) in the cis-region by an association approach and took the average across genes.

The sampling variance of 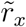 over repeated experiments can be calculated via Jackknife approach leaving one gene out at a time.

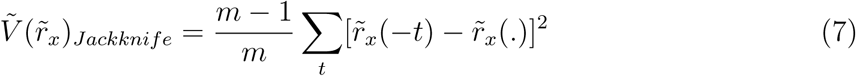

where 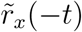 is the estimate with the *t*-th gene left out and 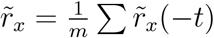. The method is derived based on eQTL data.

## 3. Results

### 3.1 DEG analysis of datasets

DEG analysis of human RNAseq and microarray datasets, comparing disease affected tissues, was performed using DESeq2 and Limma (Bioconductor packages). We used publicly available RNAseq and microarray data for brain and blood cells in PD affected individuals and controls. Such studies compare affected vs unaffected individuals to reveal differentially expressed genes (DEG) for each disease. Genes with false discovery rate (FDR) below 0.05 and a threshold of *log*_2_(1) (2-fold) increase or decrease in transcript levels was required for a gene to be accepted as a DEG. The numbers of unfiltered DEG identified were 487 in brain and 1083 in blood datasets.

We also performed cross comparative analysis to find the common significant genes between brain and blood cells. DEG common in brain and blood cells of PD affected people were identified and are summarised graphically in Figures 1 and 2. We observed that the number of common positive significant genes between brain and blood cells is 7, and their log fold changes and negative log p-values are shown in figure 1. Similarly the number of negative dysregulated significant genes between brain and blood cells is 7, and their log fold changes and negative logarithmic p-values are shown in figure 2.

**Figure 2:**
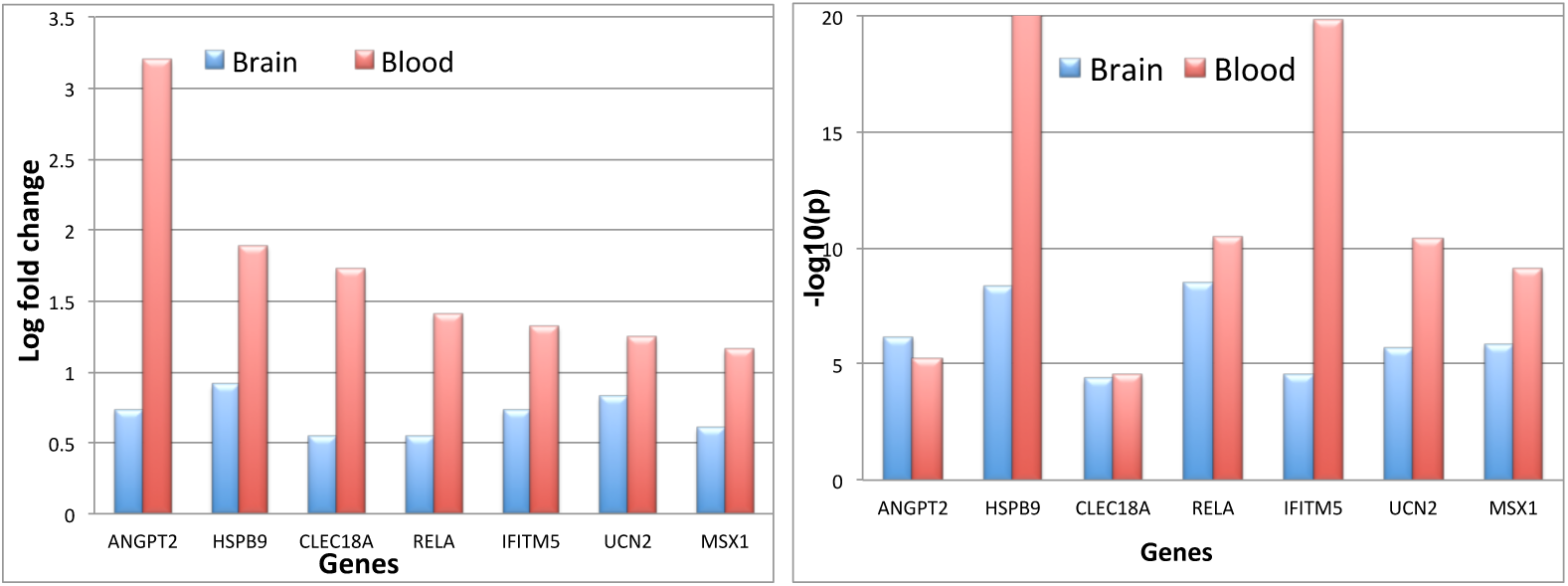
Identification of upregulated DEG that is observed inn both brain and blood. A) log fold changes of the common upregulated significant genes in brain and blood and B) Negative log of p-value of the upregulated significant genes common to brain and blood.

**Figure 3:**
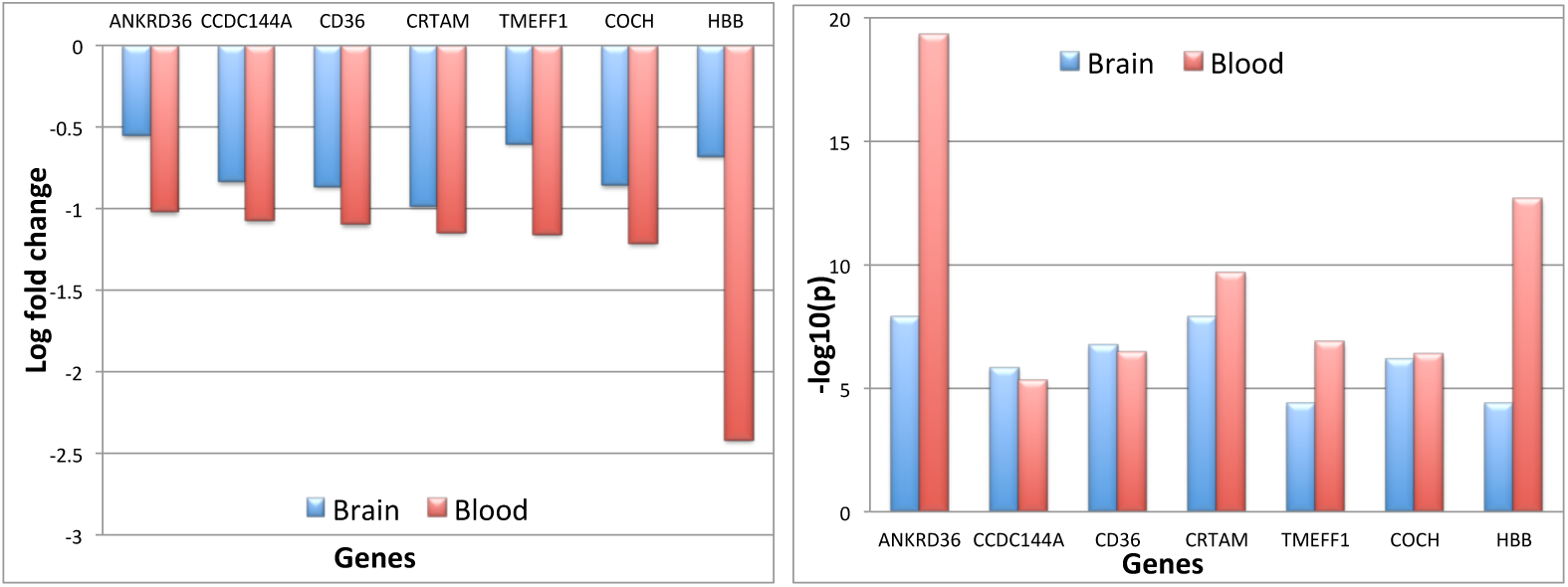
Identification downregulated DEG in both brain and blood. A) log fold changes of the common downregulated significant genes in brain and blood and B) Negative log of p-values of downregulated significant genes common to brain and blood.

### 3.2 Identifying genes expressed in blood cells that mirror those expressed in brain

eQTL databases link gene SNPs to gene expression. Few such databases have been produced, but there are databases for blood and brain cells; we used these from the GTEx database to find genes with similar genetic control of expression in the two tissues using meta-analysis approaches. We have identified 673 such blood-brain co-expressed genes (BBCG) using the correlation and meta-analysis approach as explained in the method section.

### 3.3 Identifying genes in blood that influence PD development

We selected genes whose expression or gene sequence variants (e.g., SNPs) reveal functional association with PD. This utilises curated gold-benchmark databases OMIM and GWAS catalogues. Using SNP and gene expression datasets from both brain and blood cells we identified BBCG among PD-associated genes. Similarly, using SNP and common cis-eQTL in brain and blood cells we also identified BBCG among PD-associated genes. Thus we identified 12 significant genes, C10orf32, CCDC82, COL5A2, COQ7, GPNMB, HSD17B1, KANSL1, NCKIPSD, PM20D1, SP1, FRRS1L and IL1R2 that are commonly dysregulated between blood and brain for the PD. Disease processes will alter their expression but systemic disease factors may similarly affect brain and blood cells. We used these potential PD biomarkers to identify pathways of regulation that are active in PD patients.

### 3.4 Identifying pathways in blood cells that mirror in brain

We performed pathway and gene ontology analysis on DEG sets using DAVID bioinformatics resources. For pathways we used KEGG data enrichment was determined for the identified potential signature genes for PD progression in blood cells. To combine large scale, state of the art transcriptome and proteome analyses, we then performed a regulatory analysis in order to gain further insight into the molecular pathways associated with these common genes as well as predicted links to the affected pathways. DEG and pathways were analysed using KEGG pathway database (http://www.genome.j/kegg/pathway.html) and functional annotation analysis tool DAVID (http://niaid.abcc.ncifcrf.gov) to identify overrepresented pathway groups amongst DEG sets and to group them into functional categories. Pathways deemed significantly enriched in the common DEG sets (FDR < 0.05) were reduced by manual curation to include only those with known relevance to the diseases concerned. These data are summarised on Table I. We observed a number of relevant and significant pathways including TGF-*β* and MAPK signaling pathways.

To get further insight into the identified pathways, enriched common gene sets were processed by gene ontology methods using EnrichR (http://amp.pharm.mssm.edu/Enrichr/) which identifies related biological processes and groups them into functional categories. The list of processes were also curated for those with known involvement with the diseases of interest. The cell processes and genes identified are summarised on Table II. We observed a number of significant pathways that notably included regulation of neuron death and negative regulation of neuron death.

**Table 1:** KEGG pathway analyses to identify significant pathways for the identified potential biomarker for PD that revealed among genes expressed in common by brain and blood cells. Pathway genes and pathway adjusted p-values are indicated.

| KEGG Term | Pathways | Adjusted P-value | Genes |
| --- | --- | --- | --- |
| hsa05202 | Transcriptional misregulation in cancer | 5.01E-03 | SP1;IL1R2 |
| hsa00130 | Ubiquinone and other terpenoid-quinone biosynthesis | 6.58E-03 | COQ7 |
| hsa04913 | Ovarian steroidogenesis | 2.96E-02 | HSD17B1 |
| hsa00140 | Steroid hormone biosynthesis | 3.43E-02 | HSD17B1 |
| hsa04350 | TGF-beta signaling pathway | 4.93E-02 | SP1 |
| hsa04010 | MAPK signaling pathway | 4.28E-02 | IL1R2 |
| hsa04060 | Cytokine-cytokine receptor interaction | 4.79E-02 | IL1R2 |
| hsa01100 | Metabolic pathways | 4.79E-02 | HSD17B1;COQ7 |

### 3.5 Protein-protein interaction (PPI) analysis to identify functional sub-networks

Dysregulation in a protein subnetwork may yield dysfunctional multiple protein sub-networks. Several diseases may be caused or influenced by the malfunction of a particular protein complex [34]. Thus, two or more genes are potentially related to each other through their common association in a protein-protein interaction network. Having identified genes involved in pathways and processes common to brain and blood, we sought evidence for existing sub-networks based on known PPI. Using the enriched common disease genesets, we constructed putative PPI networks using web-based visualisation resource STRING [35]. Clustering of genes was also performed by the Markov cluster algorithm (MCL) and it was notable that many of the PPI network contained genes within one cluster, indicated in red on Fig. 4. This data provides evidence that PPI sub-network exist in our enriched genesets, and confirm the presence of relevant functional pathways.

**Figure 4:**
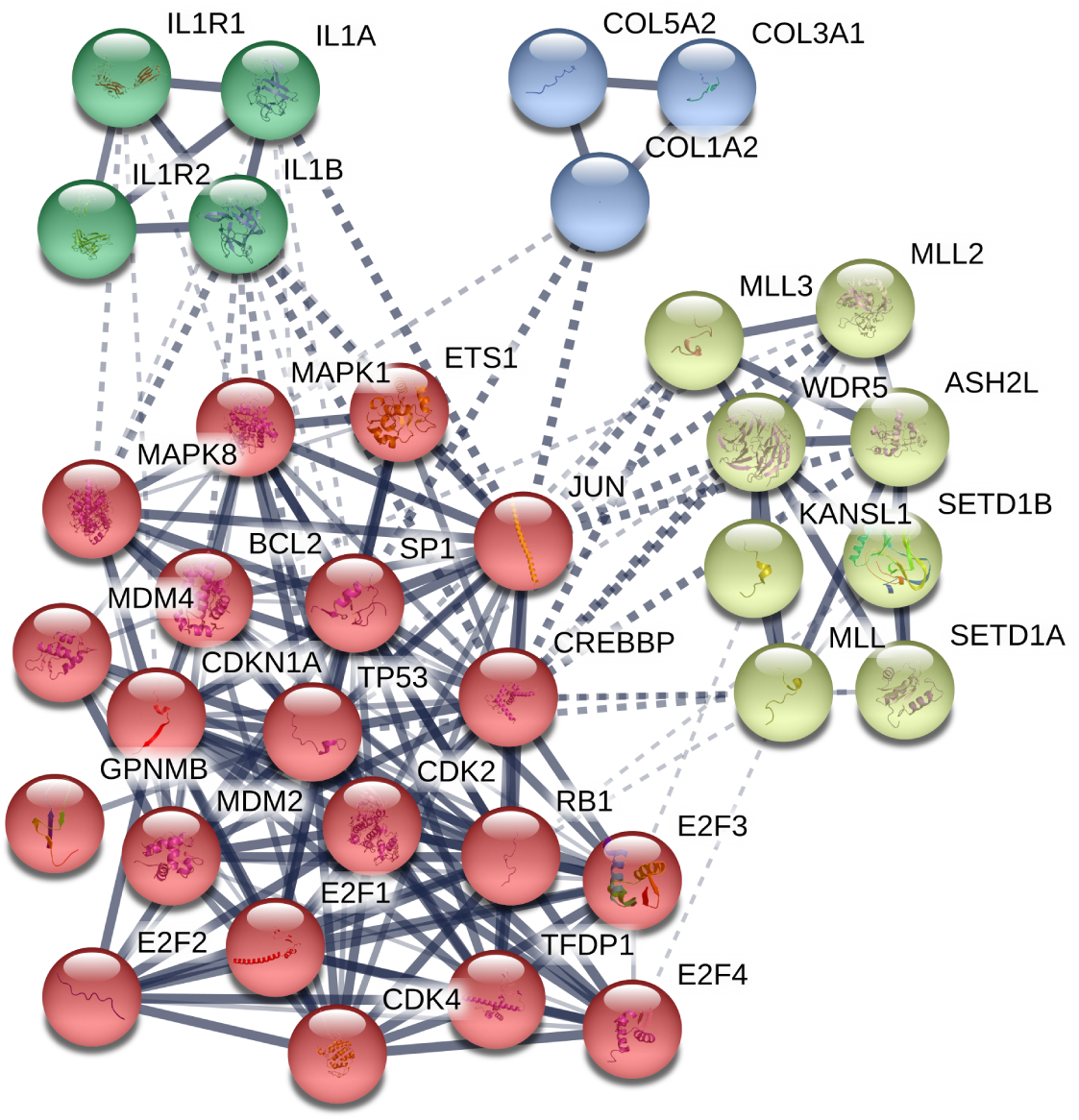
Protein-protein interaction (PPI) network of the PD. These include significant pathways common to blood and brain cells as indicated. Genes were identified by STRING software tools. Colour indicates MCL analysis clusters of proteins.

**Table 2:**
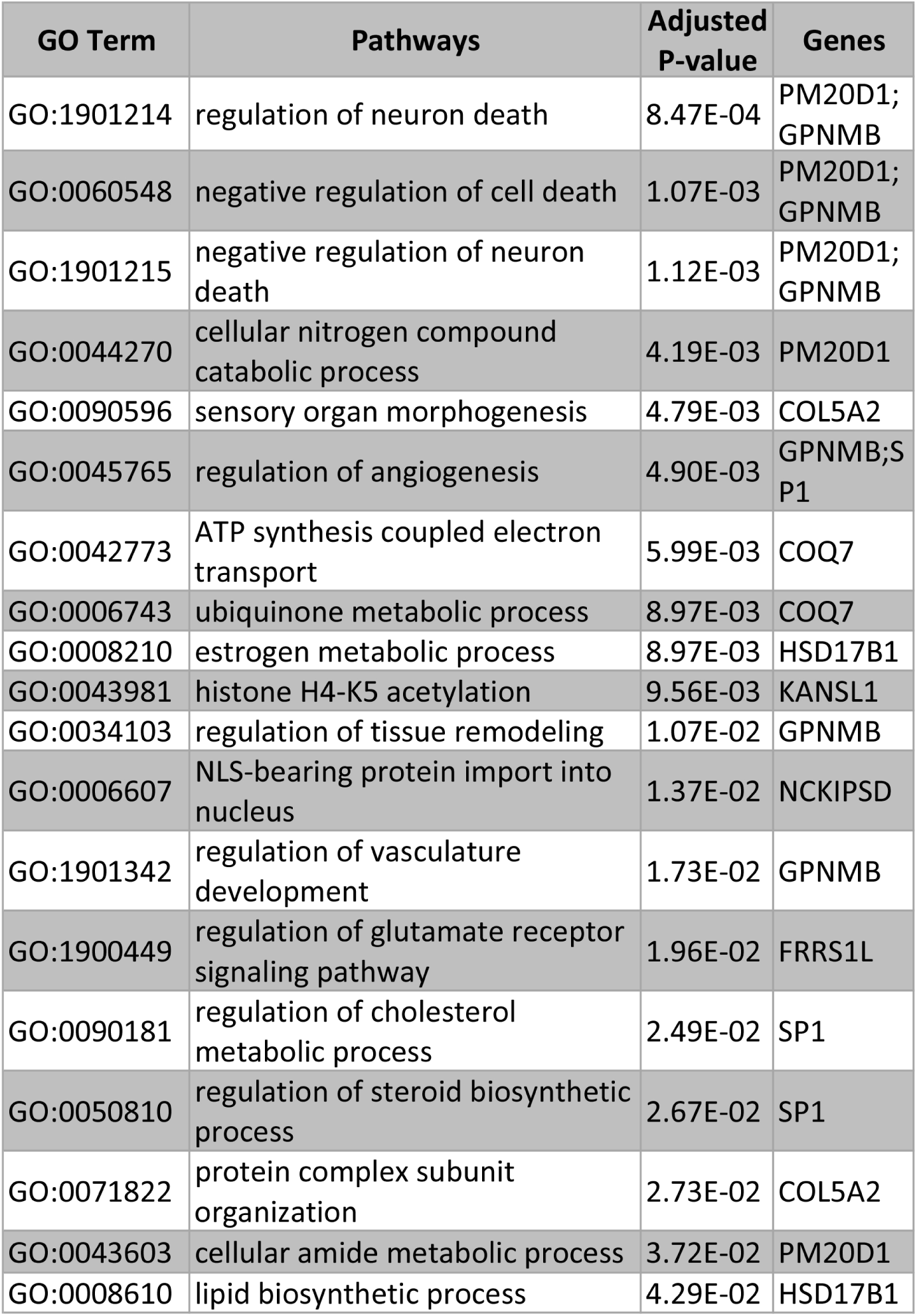
Gene ontology identification of biological processes common to brain and blood cells in PD disease. KEGG pathway-enriched gene sets were employed in gene ontology (GO) studies to identify potentially common processes. GO terms were curated to identify those relevant to brain and blood cells. Pathways genes and pathway adjusted p-values are indicated.

| GO Term | Pathways | Adjusted P-value | Genes |
| --- | --- | --- | --- |
| GO:1901214 | regulation of neuron death | 8.47E-04 | PM20D1; GPNMB |
| GO:0060548 | negative regulation of cell death | 1.07E-03 | PM20D1; GPNMB |
| GO:1901215 | negative regulation of neuron death | 1.12E-03 | PM20D1; GPNMB |
| GO:0044270 | cellular nitrogen compound catabolic process | 4.19E-03 | PM20D1 |
| GO:0090596 | sensory organ morphogenesis | 4.79E-03 | COL5A2 |
| GO:0045765 | regulation of angiogenesis | 4.90E-03 | GPNMB; SP1 |
| GO:0042773 | ATP synthesis coupled electron transport | 5.99E-03 | COQ7 |
| GO:0006743 | ubiquinone metabolic process | 8.97E-03 | COQ7 |
| GO:0008210 | estrogen metabolic process | 8.97E-03 | HSD17B1 |
| GO:0043981 | histone H4-K5 acetylation | 9.56E-03 | KANSL1 |
| GO:0034103 | regulation of tissue remodeling | 1.07E-02 | GPNMB |
| GO:0006607 | NLS-bearing protein import into nucleus | 1.37E-02 | NCKIPSD |
| GO:1901342 | regulation of vasculature development | 1.73E-02 | GPNMB |
| GO:1900449 | regulation of glutamate receptor signaling pathway | 1.96E-02 | FRRS1L |
| GO:0090181 | regulation of cholesterol metabolic process | 2.49E-02 | SP1 |
| GO:0050810 | regulation of steroid biosynthetic process | 2.67E-02 | SP1 |
| GO:0071822 | protein complex subunit organization | 2.73E-02 | COL5A2 |
| GO:0043603 | cellular amide metabolic process | 3.72E-02 | PM20D1 |
| GO:0008610 | lipid biosynthetic process | 4.29E-02 | HSD17B1 |

## 4. Discussion and Conclusions

In this study, we investigated transcriptomic evidence for intersecting pathways in brain and blood cells and tissues affected by PD disease. Employing global transcriptome analyses, we investigated in detail common gene expression profiles of PD evident in the both blood and brain cells. We investigated possible common pathways from their common patterns of gene expression, and compared these with pathways evident in validated datasets including dbGaP (see Table III) and from protein-protein interaction (PPI) data. This network-based approach identified significant common pathways influencing the both brain and blood. Moreover, we used a novel computation-based approach to identify among the common gene expression pathways that are profoundly affected by the disease processes themselves as well as by predisposing genetic and environmental factors.

In this study, we have identified potential biomarkers for PD which can be detected as transcripts in blood cells. We examined how such DEG correlate with expression of PD-associated genes to find key genes with expression dysregulated in both brain and blood cells. These are the first rationally designed candidate factors for PD evaluation using blood cell transcripts, but clinical investigations in PD patients are needed to evaluate their utility; nevertheless, their identification employed standard informatics based analytical logic and large gene expression datasets. What is novel is our approach to find disease markers that are can be employed in a tissue (blood) that is easily accessible in the clinic and does not require specialised methods of evaluation beyond gene expression analysis. The nature of these biomarkers and the the pathways they participate in may reveal new aspects of PD development and progression, particularly since these biomarkers are evident in cells outside the central nervous system, and so reflect responses to systemic factors. While the biomarkers may have practical utility it seems unlikely that they could form the basis for therapeutic developments since they do not have target tissue specificity.

**Table 3:**
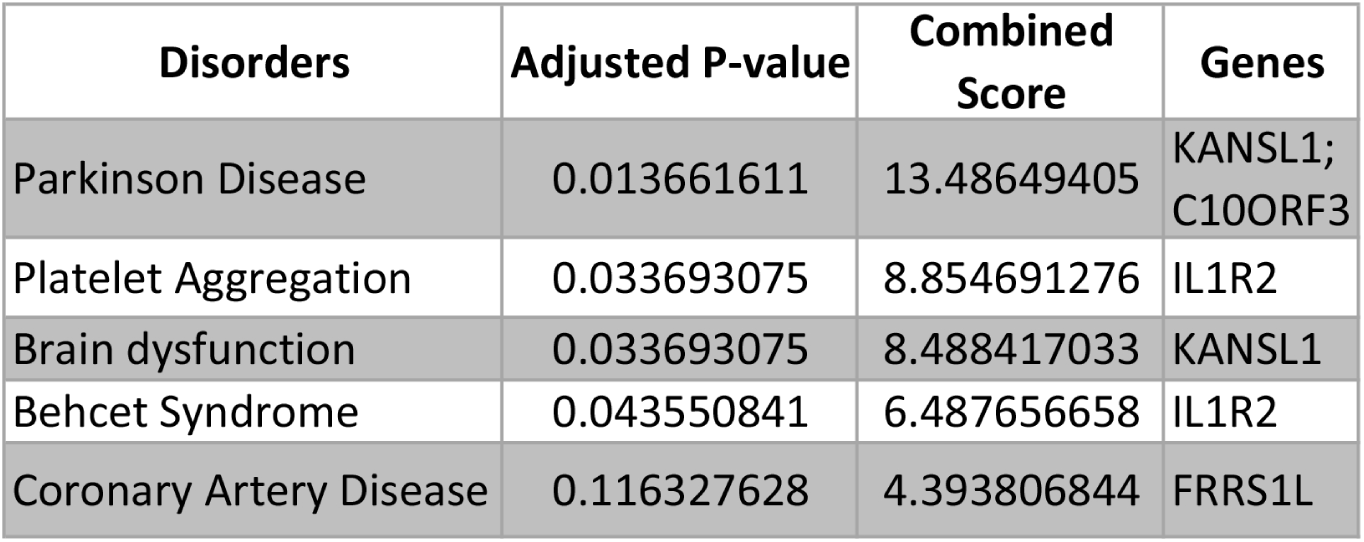
Genes significant in PD that are potential markers in blood cell for PD. These genes are also significant for in dbGaP database using single nucleotide polymorphism association with diseases.

In conclusion, our approach in identifying potential blood markers for PD has potential for diagnostic utility and could be similarly used to develop blood cell transcript-based tests for other neurodegenerative diseases that cannot yet be evaluated without detailed brain scans or surgically invasive interventions.

## Data availability

RNAseq and gene expression microarray datasets related to the blood and brain cells are available at the Gene Expression Omnibus of the National Center for Biotechnology Information (NCBI) (http://www.ncbi.nlm.nih.gov/geo/) with accession numbers *GSE*68719 and *GSE*22491. GWAS data is freely accessible in the NHGRI-EBI Catalog (https://www.ebi.ac.uk/gwas/) and eQTL data (both blood and brain) is available in the GTEx-Portal (https://gtexportal.org/home/). Pathways, ontology and PPI datasets are available respectively in the DAVID bioinformatics resources (https://david-d.ncifcrf.gov/), KEGG pathways and STRING databases string-db.org. Moreover, our incorporated gold benchmark verified datasets is freely accessible from the dbGaP (www.ncbi.nlm.nih.gov/gap).

